## Supplementary Materials for "Invadopodia enable cooperative invasion and metastasis of breast cancer cells"

**This PDF file includes:**

Figures S1 to S12

**Other Supplementary Materials for this manuscript include the following:**

Movies S1 to S10

Source data

**Figure S1**

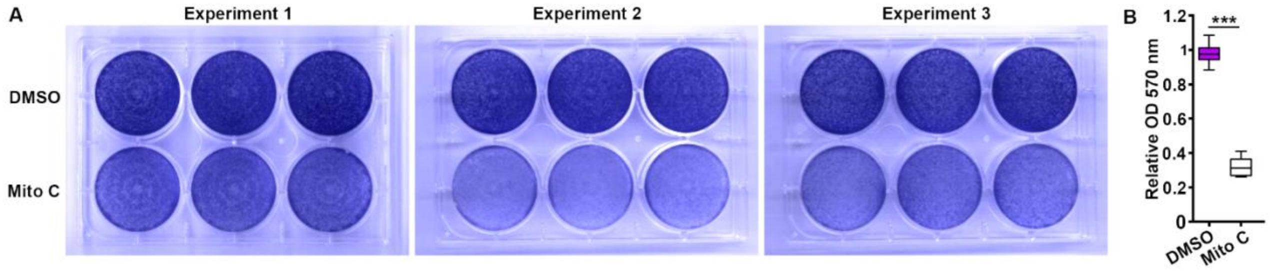


**Figure S1.** (**A**) Crystal violet staining of 4T1 cells after 2 days of treatment with DMSO (top wells) or mitomycin C (Mito C, bottom wells). Three independent experiments were conducted. (**B**) Relative optical density (OD) at 570 nm of cells in DMSO control (magenta box) and mitomycin C-treated wells (white box) from (A). $P$=2.54x$10^{-12}$, by the t-test.

**Figure S2**

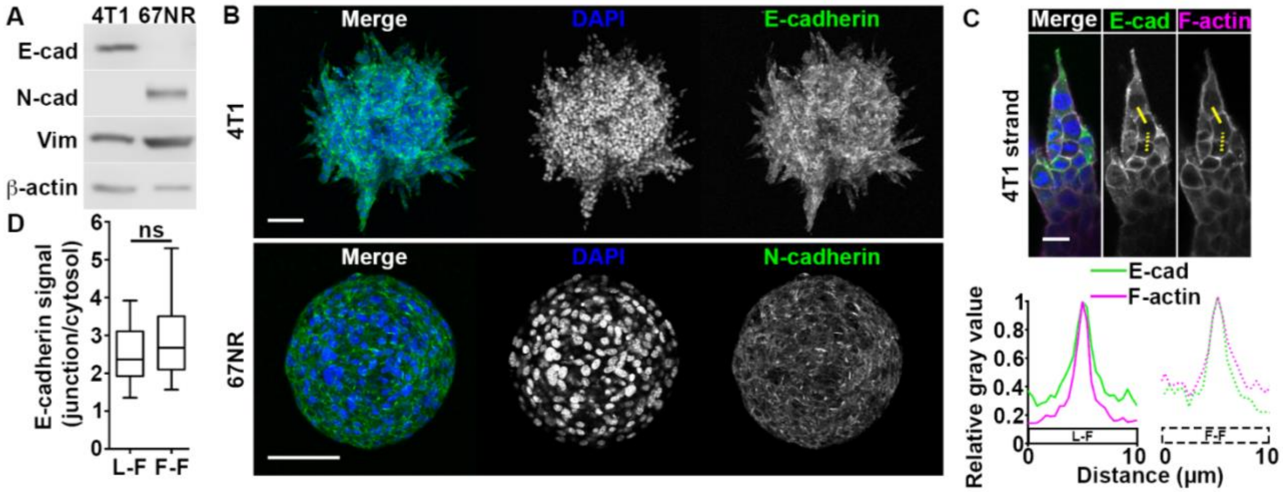


**Figure S2**. (**A**) E/N-cadherin (E/N-cad) and vimentin (vim) expression levels in 4T1 and 67NR cells. (**B**) 4T1 and 67NR spheroids at day 2 post-embedding in a 3D collagen I matrix, immunolabeled for E/N-cadherin (green) and stained with DAPI (blue). Scale bars: 100 μm. (**C**) 4T1 strand at day 2 post-embedding. E-cadherin (E-cad, green), F-actin (phalloidin, magenta) and nuclei (DAPI, blue) were stained. Bottom panels show the E-cadherin (E-cad, green) and F-actin (magenta) signals along the solid (leader cell-follower cell junction (L-F)) and dashed (follower cell-follower cell junction (F-F)) yellow lines. Scale bar: 20 μm. (**D**) Ratio of E-cadherin signal at L- F and F-F cell junctions over cytosol from (C).

**Figure S3**

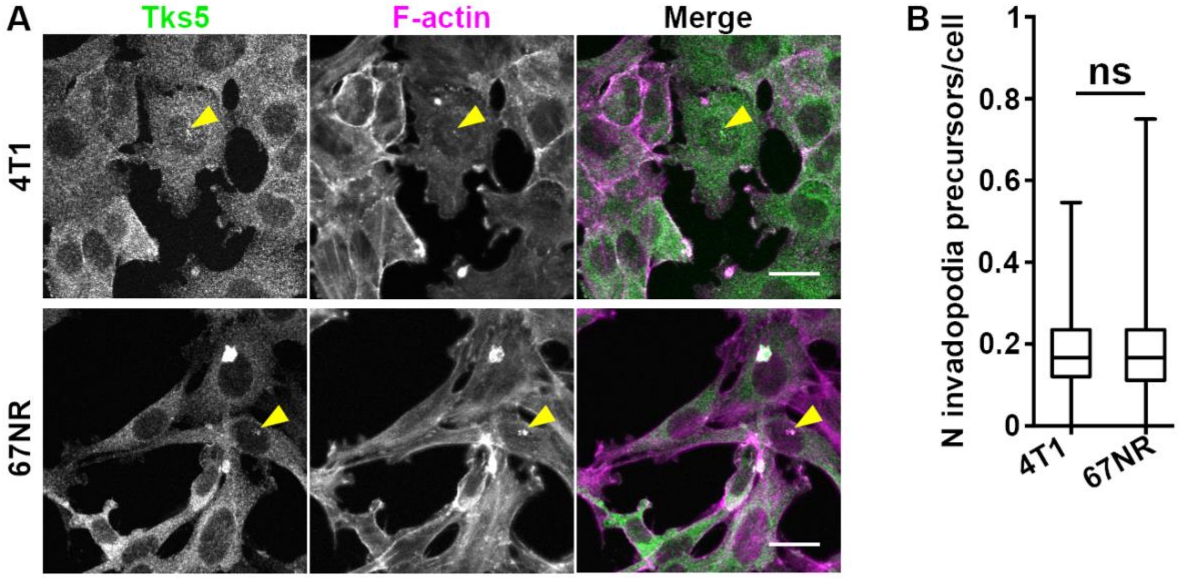


**Figure S3.** (**A**) 4T1 (top) and 67NR (bottom) cells cultured on fluorescent gelatin (not shown). Tks5 (green) and F-actin (phalloidin, magenta) were stained. Yellow arrowheads indicate invadopodia precursors. Scale bars: 20 µm. (**B**) Number of invadopodia precursors (Tks5 + F-actin) per cell from (A).

### Figure S4

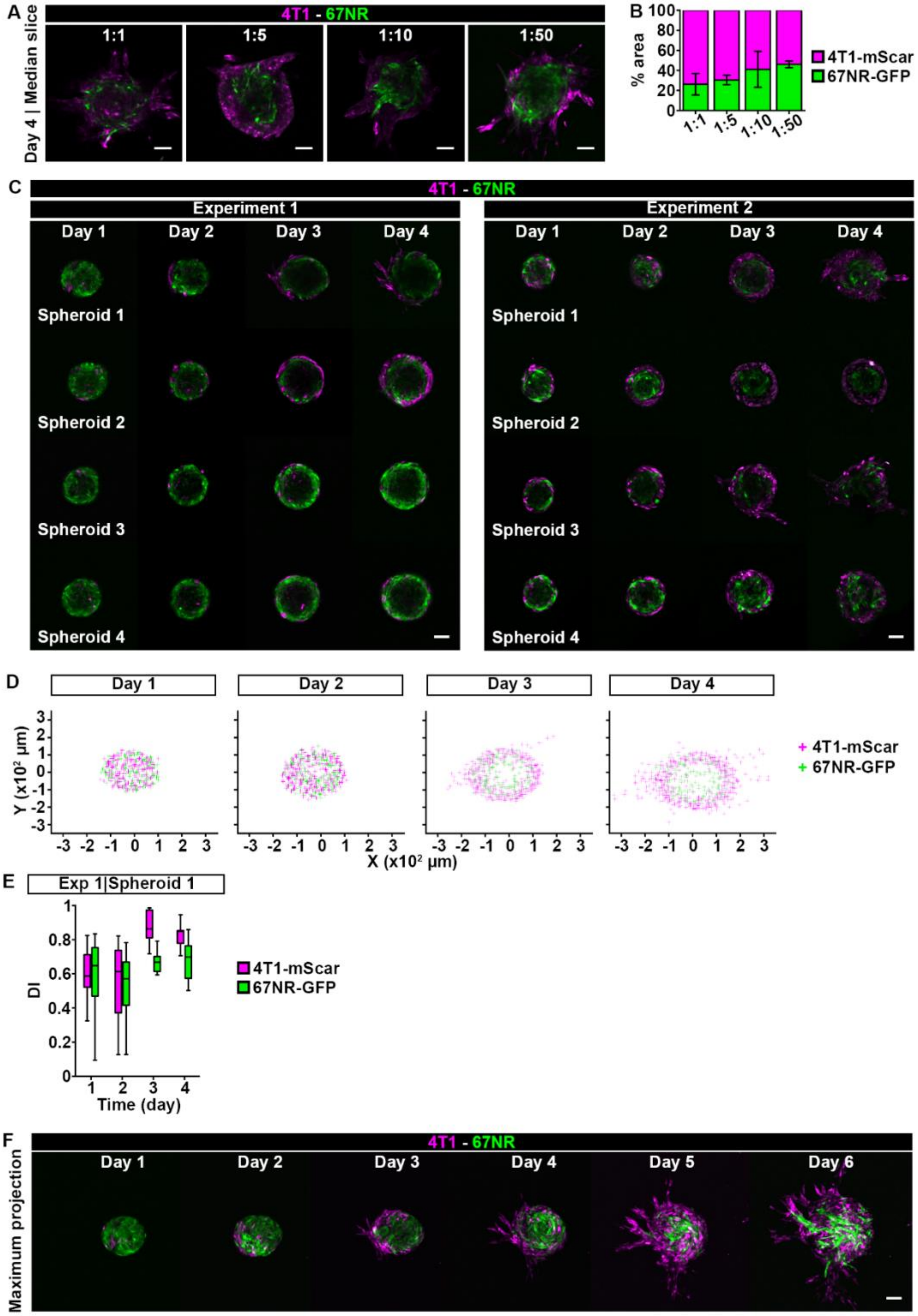
A
4T1 - 67NR
Day 4 | Median slice
1:1
1:5
1:10
1:50
B
% area
4T1-mScar
67NR-GFP
C
Experiment 1
Experiment 2
Day 1
Day 2
Day 3
Day 4
Spheroid 1
Spheroid 2
Spheroid 3
Spheroid 4
D
Y (x10² µm)
X (x10² µm)
E
Exp 1|Spheroid 1
DI
Time (day)
F
Maximum projection
Day 5
Day 6

**Figure S4. (A)** Mixed 4T1-mScarlet:67NR-GFP spheroids on day 4 post-embedding. Spheroids were made with varying ratios of 4T1-mScarlet (magenta) to 67NR-GFP (green) cells, as indicated. Scale bars: 100 µm. **(B)** Percent area occupied by 4T1-mScarlet (magenta bars) and 67NR-GFP (green bars) cells in the mixed spheroids from (A). A stacked bar graph with mean and standard error is shown. **(C)** All the mixed spheroids used for Fig. 2B and 2D. Scale bars: 100 µm. **(D)** Relative coordinates of 4T1-mScarlet (magenta crosses) and 67NR-GFP (green crosses) cells from all mixed spheroids presented in (A) and Fig. S3C, including cells in invasion strands. **(E)** Distance Index (DI) for 4T1-mScarlet (magenta boxes) and 67NR-GFP (green boxes) cells from spheroid 1 in experiment 1 from (C). **(F)** Mixed spheroid at a 1:50 ratio of 4T1-mScarlet to 67NR-GFP cells, day 1-6 post-embedding. See Fig. 2A. Scale bar: 100 µm.

**Figure S5**

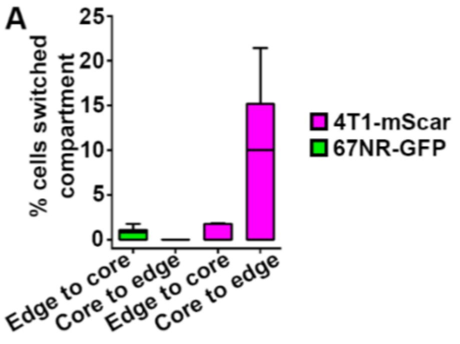


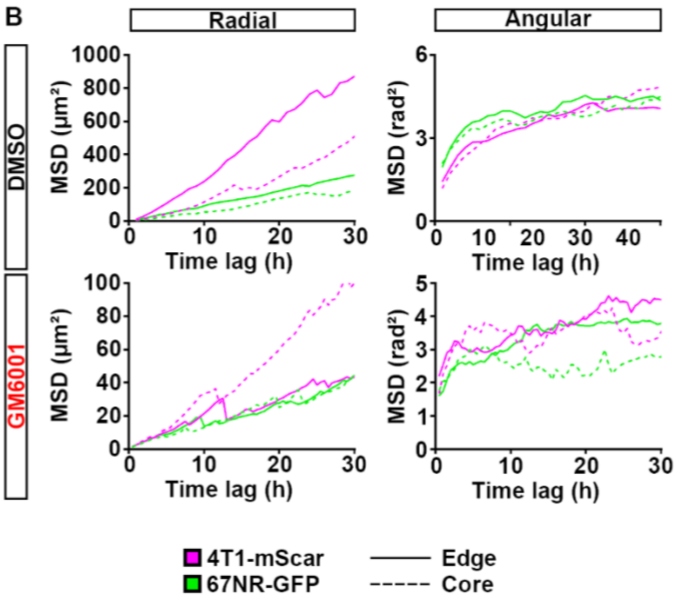


C

DMSO
GM6001

**Radial**

| | $\alpha_r$ | $\Gamma_r$ (µm²) | R |
|---|---|---|---|
| 4T1-mScar \| Edge | 1.2<br>0.94 | 15<br>1.2 | 0.996<br>0.983 |
| 4T1-mScar \| Core | 1.4<br>1.4 | 4.0<br>0.5 | 0.995<br>0.987 |
| 67NR-GFP \| Edge | 1.0<br>0.88 | 9.1<br>1.3 | 0.992<br>0.931 |
| 67NR-GFP \| Core | 1.1<br>0.69 | 4.4<br>1.7 | 0.990<br>0.951 |

**Angular**

| | $\alpha_\phi$ | $\Gamma_\phi$ (rad²) | R |
|---|---|---|---|
| 4T1-mScar \| Edge | 0.3<br>0.08 | 1.6<br>2.5 | 0.962<br>0.542 |
| 4T1-mScar \| Core | 0.5<br>0.2 | 1.2<br>2.1 | 0.997<br>0.819 |
| 67NR-GFP \| Edge | 0.3<br>0.2 | 2.2<br>1.7 | 0.964<br>0.926 |
| 67NR-GFP \| Core | 0.2<br>0.1 | 2.2<br>2.0 | 0.964<br>0.575 |

D

DMSO
GM6001

**Radial**

| | $D_{eff}$(3 h) (µm²/s) |
|---|---|
| 4T1-mScar \| Edge | 12<br>0.53 |
| 4T1-mScar \| Core | 5.5<br>0.63 |
| 67NR-GFP \| Edge | 4.6<br>0.48 |
| 67NR-GFP \| Core | 2.7<br>0.40 |

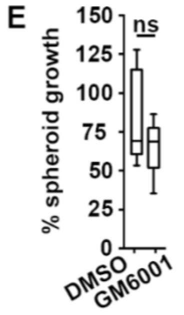


**Figure S5. (A)** Percentage of 4T1-mScarlet (magenta) and 67NR-GFP (green) cells that switched compartments (from edge to core or *vice versa*), see Fig. 2H. **(B)** Mean square displacements (MSDs) for 4T1-mScarlet (magenta) and 67NR-GFP (green) cells, see Fig. 2H. MSDs were calculated in the radial (r, left panels) and angular ($\phi$, right panels) directions of the polar coordinate system, for edge (solid lines) and core (dashed lines) cells in spheroids treated with DMSO (top panels) and GM6001 (bottom panels), respectively. **(C)** Slope ($\alpha_{r,\phi}$), intercept ($\Gamma_{r,\phi}$) and R (goodness of fit) values for the power law fit to the MSD data from (B). See Materials and Methods for details. Values for DMSO-treated spheroids are in black and for GM6001-treated spheroids in red. **(D)** Effective diffusion coefficient ($D_{eff}$) in the radial direction (see Materials and Methods) calculated at 3 h based on the values from (B). **(E)** Spheroid growth after 4 days of treatment with GM6001 or DMSO (control), see Figs. 2A, 2E.

### Figure S6

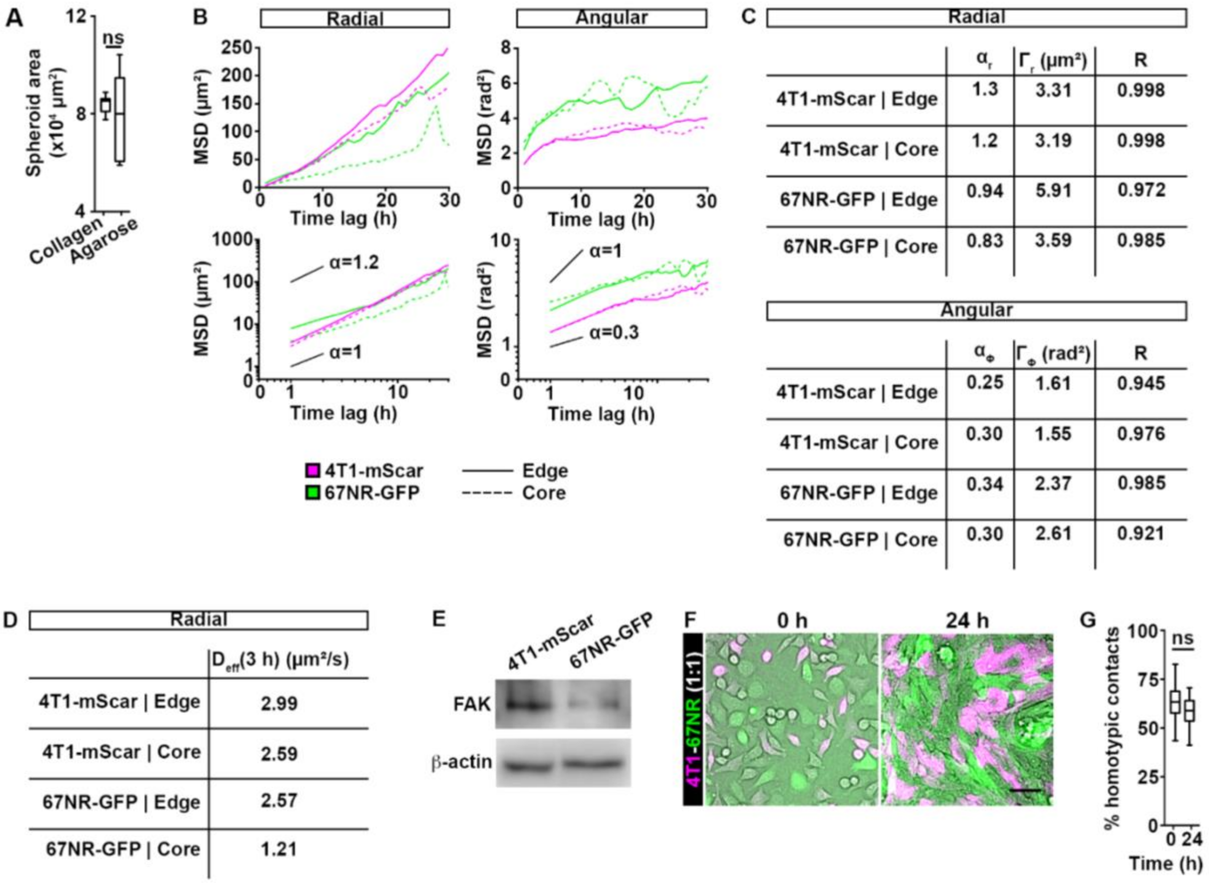


**Figure S6.** (**A**) Spheroid area at day 3 post-embedding in a 3D collagen I or agarose matrix, see Figs. 2A, 3A. (**B**) MSDs for 4T1-mScarlet (magenta) and 67NR-GFP (green) cells, see Fig. 3C. MSDs were calculated in the radial (r, left top panel) and angular (ϕ, right top panel) directions of the polar coordinate system, for edge (solid lines) and core (dashed lines) cells in spheroids embedded in agarose. The bottom panels correspond to the log-log plots of the top panels, indicating the fundamentally different mechanisms of transport, *i.e.* sub-diffusive for $\alpha<1$, diffusive for $\alpha=1$, and super-diffusive for $\alpha>1$. Solid lines serve as guides, indicating the average slopes (α values) corresponding to these different motility modalities. (**C**) Slope ($\alpha_{r,\phi}$), intercept ($\Gamma_{r,\phi}$) and R (goodness of fit) values for the power law fit to the MSD data from (B). See Materials and Methods for details. (**D**) Effective diffusion coefficient ($D_{eff}$) in the radial direction calculated at 3 h, based on the values from (B). (**E**) Western blot analysis of phosphorylated FAK (pFAK) and FAK expression in 4T1-mScarlet and 67NR-GFP cells. β-actin was used as a loading control. (**F**) 4T1-mScarlet (magenta) and 67NR-GFP (green) cells at 0 h (left) and 24 h (right) post-plating on gelatin, at a 1:1 ratio. Scale bar: 50 μm. (**G**) Percentage of homotypic contacts in cells from (F).

**Figure S7**

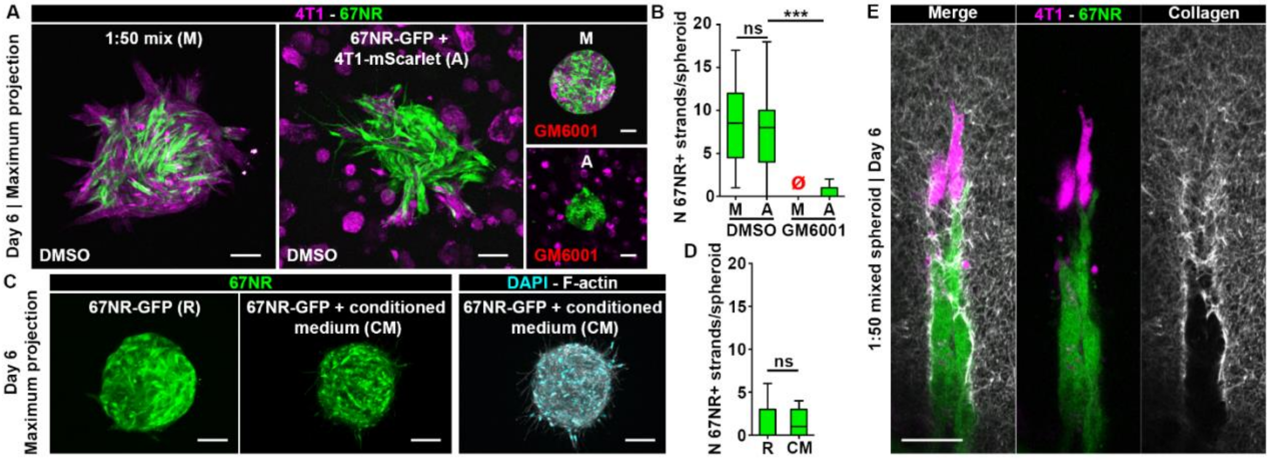


**Figure S7.** (**A**) Mixed spheroids of 4T1-mScarlet and 67NR-GFP at a 1:50 ratio, ***M***, or spheroids of 67NR-GFP cells with 4T1-mScarlet cells added in the collagen, ***A***. Spheroids were treated from day 0 with GM6001 (right panels) or DMSO control (left panels) and imaged at day 6. Scale bars: 100 µm. (**B**) Number of strands containing 67NR cells (67NR+) per spheroid from (A). The red empty symbols indicate zero values. $P$=8.90x$10^{-6}$, by the Wilcoxon rank sum test. (**C**) 67NR-GFP spheroids treated with regular medium, ***R***, or conditioned medium, ***CM***, collected from 4T1-mScarlet cells grown on gelatin. Nuclei (DAPI, cyan) and F-actin (phalloidin, white) were stained. Scale bars: 100 µm. (**D**) Number of strands containing 67NR cells (67NR+) per spheroid from (C). (**E**) Strand from a mixed spheroid, including collagen labeling (white). Scale bar: 50 µm.

**Figure S8**

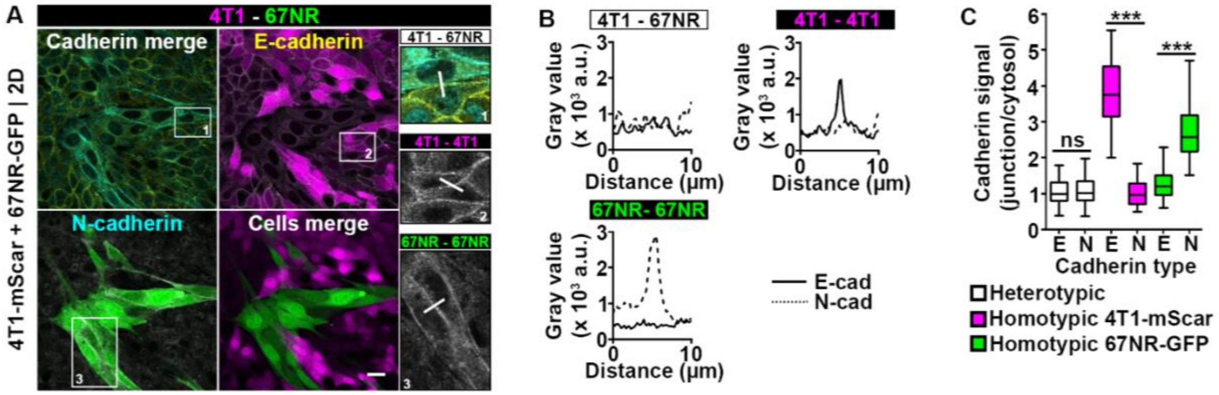


**Figure S8**. (**A**) 4T1-mScarlet and 67NR-GFP cells in 2D, immunolabeled for E/N-cadherin (yellow/cyan). The insets show a 2X zoom-in of the boxed areas 1-3. Scale bar: 20 µm. (**B**) Relative E/N-cadherin (solid/dashed line) signals along the lines in the insets 1-3 from (A). (**C**) E/N-cadherin signal of the junction over cytosol for homotypic and heterotypic junctions from (B). $P<2.20 \times 10^{-16}$ and $4.00 \times 10^{-14}$, by the Wilcoxon rank sum test.

**Figure S9**

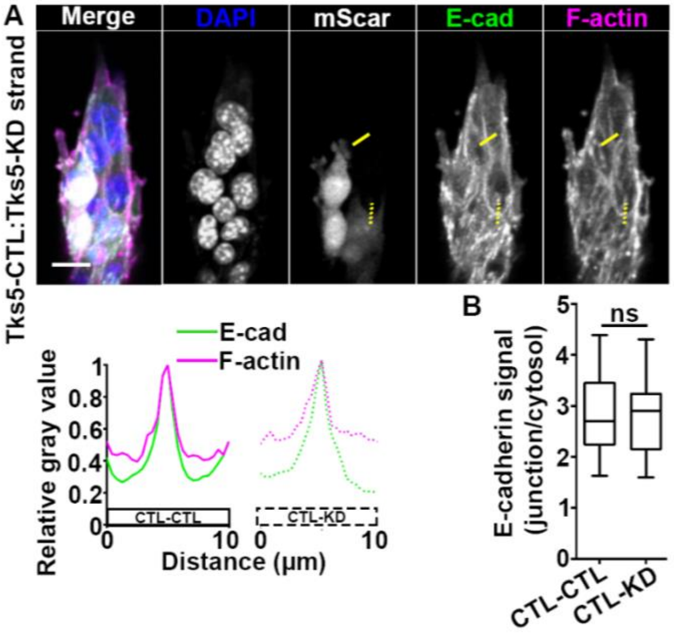


**Figure S9.** (**A**) Mixed Tks5-CTL:Tks5-KD strand, day 2 post-embedding. The spheroid was immunolabeled for E-cadherin (E-cad, green), and F-actin (phalloidin, magenta) and nuclei (DAPI, blue) were stained. Bottom panels show the relative E-cadherin (green) and F-actin (magenta) signals along the solid (CTL-CTL junction) and dashed (CTL-KD junction) yellow lines. Scale bar: 20 µm. (**B**) Relative E-cadherin signal at CTL-CTL and CTL-KD junctions over cytosol from (A).

**Figure S10**

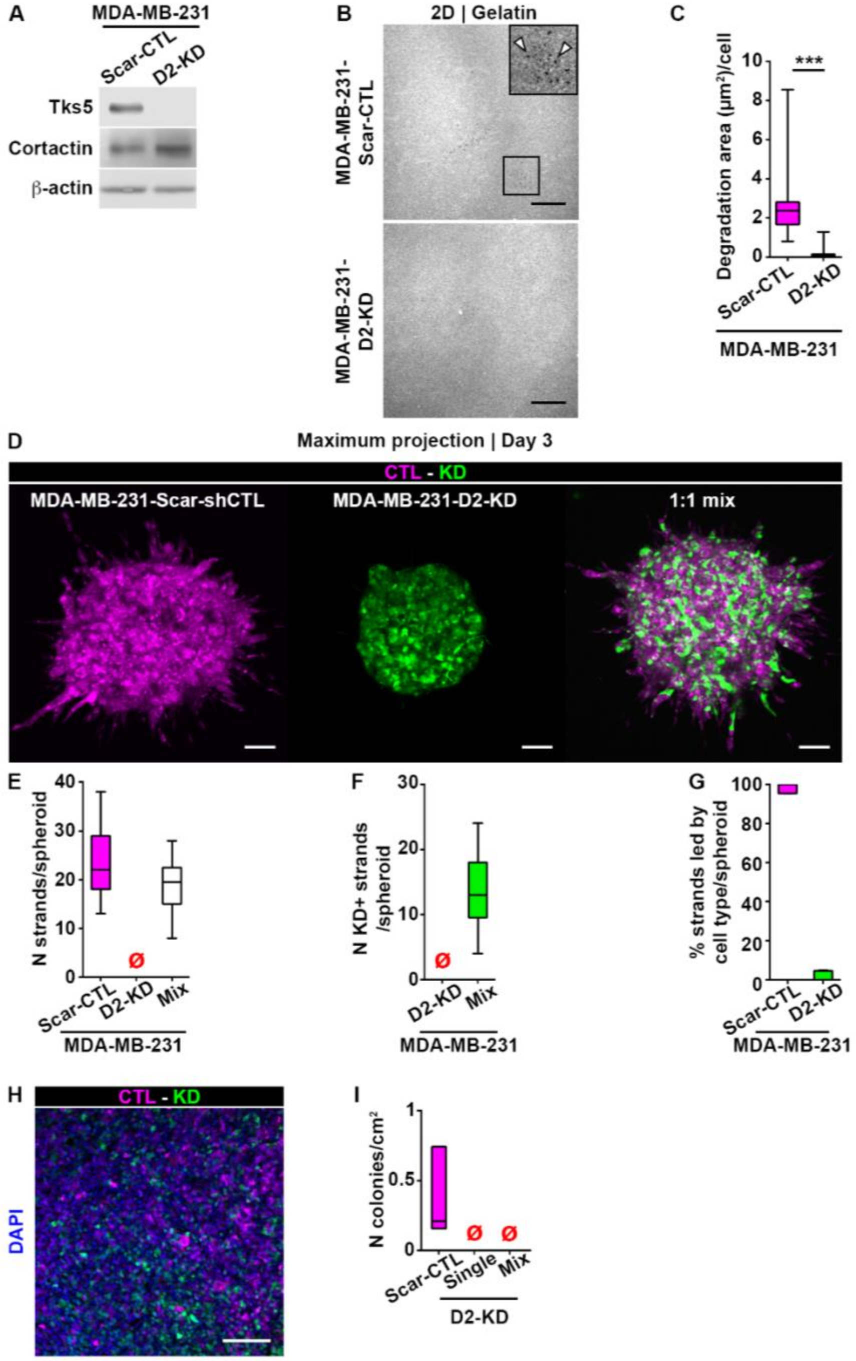


**Figure S10**. (**A**) Tks5 and cortactin expression in MDA-MB-231-mScarlet-CTL (Scar-CTL) and MDA-MB-231-Dendra2-hTks5 KD (D2-KD) cells. β-actin was used as a loading control. (**B, C**) Gelatin degradation for Scar-CTL (top panel, magenta box) and D2-KD (bottom panel, white box), 18 h after plating. The inset shows a 2X zoom-in of the boxed area; arrowheads indicate degradation holes. Scale bars: 20 µm. $P<2.20\times10^{-16}$, by the Wilcoxon rank sum test. (**D-G**) Single or mixed Scar-CTL and D2-KD spheroids, at a 1:1 ratio, day 3 post-embedding. Number of strands per spheroid (E), number of strands containing D2-KD cells (KD+) per spheroid (F) and percentage of strands led by Scar-CTL or D2-KD cells (G) from spheroids in (D). The red empty symbols indicate zero values. Scale bars: 100 µm. (**H**) Section from a mixed Scar-CTL (magenta) and D2-KD (green) tumor, with nuclei labeled with DAPI. Scale bar: 50 µm. (**I**) Number of lung colonies per $cm^2$ for mice inoculated with Scar-CTL, D2-KD or a mixture of Scar-CTL and D2-KD cells. The red empty symbols indicate zero values.

**Figure S11**

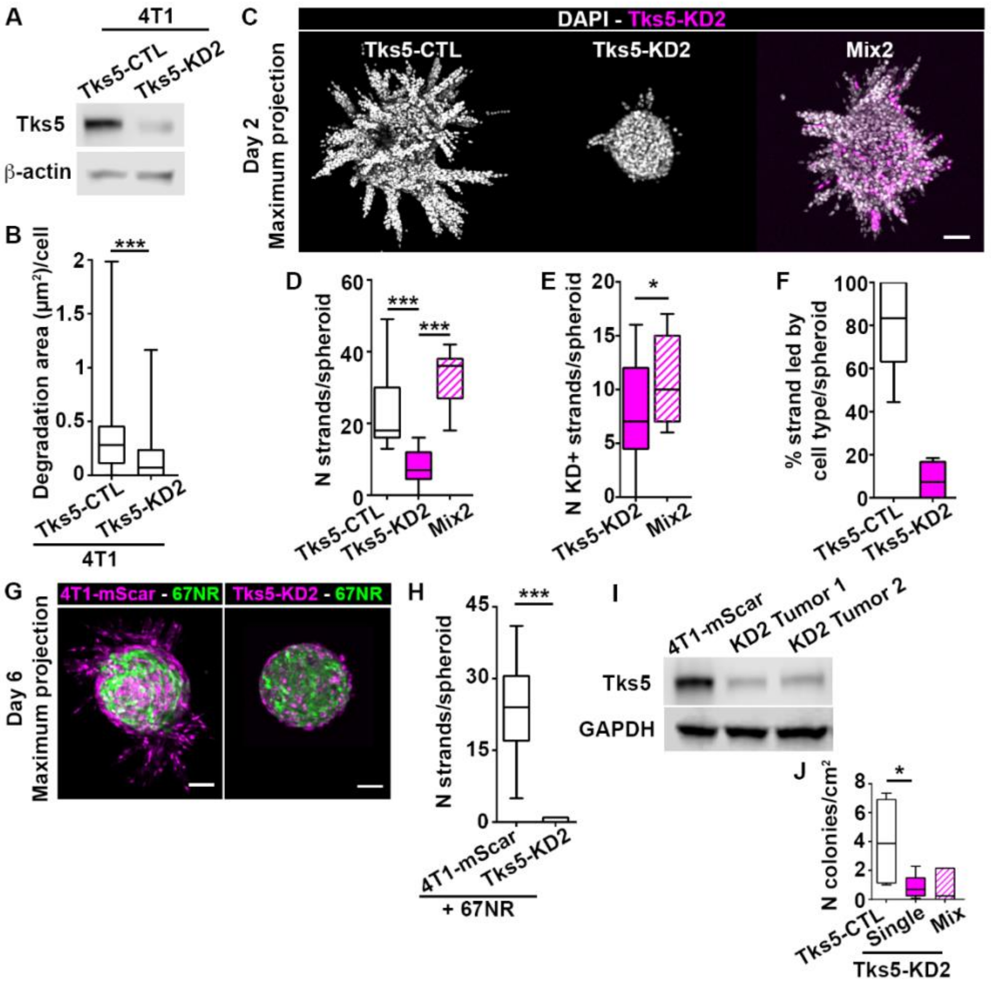


**Figure S11.** (**A**) Tks5 expression in Tks5-CTL and Tks5-KD2 cells. Knockdown efficiency is 81.6% (**B**) Degradation area (μm²) per cell for Tks5-CTL and Tks5-KD2 cells plated on gelatin. $P$=7.12×10$^{-7}$, by the Wilcoxon rank sum test. (**C**) Day 2 images of spheroids made with Tks5-CTL, -KD2 or a mixture (1:1 ratio) of Tks5-CTL and -KD2 cells. Scale bar: 100 μm. (**D-F**) Number of strands per spheroid (D), number of strands containing Tks5-KD2 cells (E), and the percentage of strands led by Tks5-CTL and -KD2 cells (F) in spheroids from (C). $P$=4.26×10$^{-6}$ and $P$=4.78×10$^{-5}$, by the Wilcoxon rank sum test in (E). $P$=0.0404, by the t-test in (F). (**G**) Day 6 images of mixed spheroids made with 67NR-GFP and 4T1-mScarlet or Tks5-KD2 cells, at a 1:50 ratio. Scale bars: 100 μm. (**H**) Number of strands per spheroid from (G). P=1.40×10$^{-4}$, by the Wilcoxon rank sum test. (**I**) Tks5 expression in 4T1-mScarlet (4T1-mScar) or Tks5-KD2 tumors. GAPDH was used as a loading control. (**J**) Number of lung colonies per cm² for mice inoculated with Tks5-CTL, Tks5-KD2 or a mixture of Tks5-CTL and Tks5-KD2 cells. $P$=0.044, by the t-test.

**Figure S12**

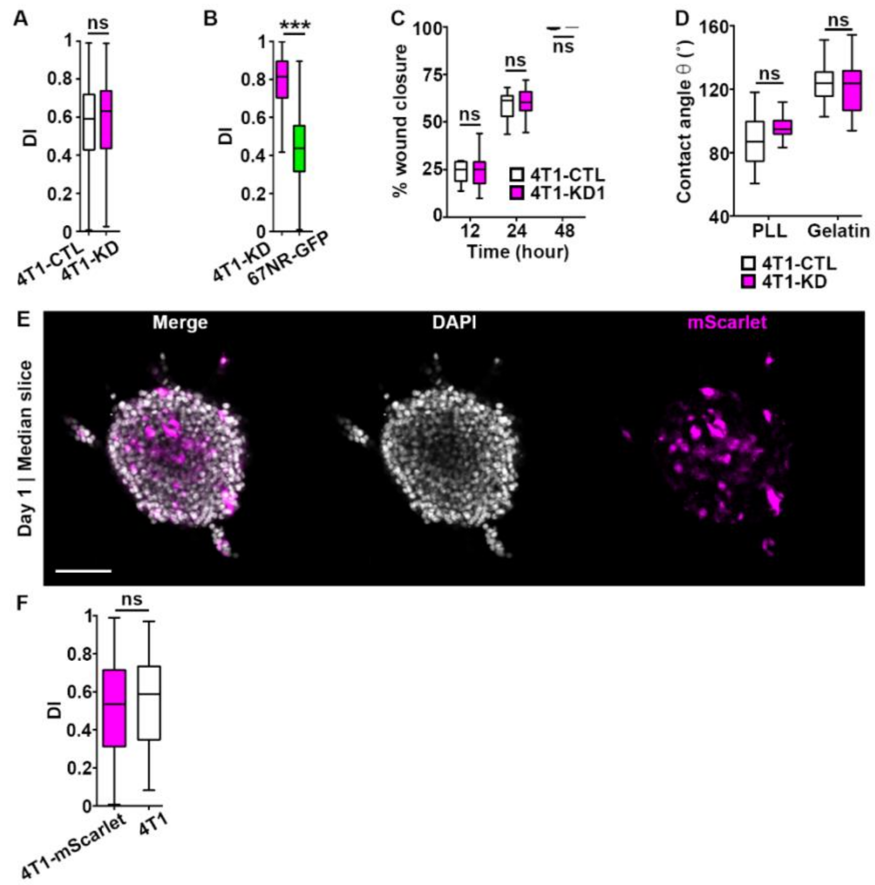


**Figure S12.** (**A**) DI for Tks5-CTL (white box) and Tks5-KD (magenta box) cells from spheroids in Fig. 6D. (**B**) DI for Tks5-KD and 67NR-GFP (green box) cells from spheroids in Fig. 6H. $P<2.20x10^{-16}$, by the Wilcoxon rank sum test. (**C**) Wound closure over time for Tks5-CTL and Tks5-KD. (**D**) Contact angle θ between Tks5-CTL or Tks5-KD on poly-L-lysine (PLL) or gelatin, 5 h post-plating. (**E, F**) Day 1 image (E) and DI (F) for 4T1-mScarlet (magenta) and wild type 4T1 (white) mixed spheroid (1:1 ratio). Scale bar: 100 µm.

### Supplementary Movies

**Movie S1.**

Time lapse of a 4T1 (top panel) and a 67NR (bottom panel) monolayer in the scratch assay. Time is in hh:mm. Scale bar: 100 µm.

**Movie S2.**

Confocal z stack of a mixed 4T1-mScarlet (magenta) and 67NR-GFP (green) spheroid grown in a 3D collagen I matrix for 3 days. Scale bar: 100 µm.

**Movie S3.**

Time lapse of 4T1-mScarlet cells in a mixed spheroid embedded in collagen I. Spheroids were treated from day 0 with DMSO control. 67NR-GFP cells are not shown to ease visualization. The left panel shows all trajectories, the middle panel shows a representative core trajectory (from edge to core) and the right panel shows a representative edge trajectory (from edge to edge). Time is in hh:mm. Scale bar: 100 µm.

**Movie S4.**

Time lapse of 67NR-GFP cells in a mixed spheroid embedded in collagen I. Spheroids were treated from day 0 with DMSO control. 4T1-mScarlet cells are not shown to ease visualization. The left panel shows all trajectories, the middle panel shows a representative core trajectory (from core to edge) and the right panel shows a representative edge trajectory (from edge to edge). Time is in hh:mm. Scale bar: 100 µm.

**Movie S5.**

Time lapse of 4T1-mScarlet cells in a mixed spheroid embedded in collagen I. Spheroids were treated from day 0 with GM6001. 67NR-GFP cells are not shown to ease visualization. The left panel shows edge trajectories and the right panel shows edge trajectories. Time is in hh:mm. Scale bar: 100 µm.

**Movie S6.**

Time lapse of 67NR-GFP cells in a mixed spheroid embedded in collagen I. Spheroids were treated from day 0 with GM6001. 4T1-mScarlet cells are not shown to ease visualization. The left panel shows edge trajectories and the right panel shows edge trajectories. Time is in hh:mm. Scale bar: 100 µm.

**Movie S7.**

Time lapse of 4T1-mScarlet cells in a mixed spheroid embedded in agarose. 67NR-GFP cells are not shown to ease visualization. Representative trajectories are shown. Time is in hh:mm. Scale bar: 100 µm.

**Movie S8.**

Time lapse of 4T1-mScarlet cells in a mixed spheroid embedded in agarose. 67NR-GFP cells are not shown to ease visualization. Representative edge trajectories are shown to illustrate the inward movement of cells. Time is in hh:mm. Scale bar: 100 µm.

**Movie S9.**
Time lapse of 4T1-mScarlet cells in a mixed spheroid embedded in agarose. 67NR-GFP cells are not shown to ease visualization. A representative core trajectory is shown to illustrate a cell reaching then leaving the edge compartment. Time is in hh:mm. Scale bar: 100 µm.

**Movie S10.**
Time lapse of 4T1-mScarlet (gray) cells at the gelatin/poly-L-lysine interface (red line). A cell that crossed the interface and migrated back to the gelatin layer is indicated with a yellow arrowhead. Time is in hh:mm. Scale bar: 50 µm.
